# Reconstitution of the Human tRNA Splicing Endonuclease Complex: insight into the regulation of pre-tRNA cleavage

**DOI:** 10.1101/2019.12.16.878546

**Authors:** Cassandra K. Hayne, Casey A. Schmidt, A. Gregory Matera, Robin E. Stanley

## Abstract

The splicing of tRNA introns is a critical step in pre-tRNA maturation. In archaea and eukaryotes, tRNA intron removal is catalyzed by the tRNA splicing endonuclease (TSEN) complex. Eukaryotic TSEN is comprised of four core subunits (TSEN54, TSEN2, TSEN34, and TSEN15). The human TSEN complex additionally co-purifies with the polynucleotide kinase CLP1; however, CLP1’s role in tRNA splicing remains unclear. Mutations in genes encoding all four TSEN subunits, as well as CLP1, are known to cause neurodegenerative disorders, yet the mechanisms underlying the pathogenesis of these disorders are unknown. Here, we developed a recombinant system that produces active TSEN complex. Co-expression of all four TSEN subunits is required for efficient formation and function of the complex. We show that human CLP1 associates with the active TSEN complex, but is not required for tRNA intron cleavage *in vitro*. Moreover, RNAi knockdown of the *Drosophila* CLP1 orthologue, cbc, promotes biogenesis of mature tRNAs and circularized tRNA introns (tricRNAs) *in vivo*. Collectively, these and other findings suggest that CLP1/cbc plays a regulatory role in tRNA splicing by serving as a negative modulator of the direct tRNA ligation pathway in animal cells.

## INTRODUCTION

Transfer RNAs (tRNAs) play a critical role in the translation of mRNA into proteins. Eukaryotic tRNA genes are transcribed by RNA Pol III and undergo a series of post-transcriptional processing and modification steps prior to reaching their mature and functional state (1–6). A subset of tRNA genes contain introns that must be removed during tRNA maturation. Intron-containing tRNA genes are found across all three domains of life (7–9). In bacteria, tRNA introns are self-spliced (8), whereas archaea and eukaryotes utilize a tRNA splicing endonuclease (TSEN) complex to catalyze the removal of introns (1,8,10–12). Archaeal tRNA splicing endonucleases have been sorted into four classes, based upon the subunit organization, including: homotetramers (α_4_), homodimers (α_2_ or ε_2_), or dimers of dimers ((αβ)_2_) (1). Conversely, the eukaryotic tRNA splicing endonuclease (TSEN) complexes are composed of four individual subunits (TSEN2, TSEN34, TSEN54, and TSEN15) (13,14) (Figure 1). TSEN2 and TSEN34 are metal-ion independent nucleases that cleave the 5’ and 3’ splice sites respectively and generate the 5’ exon with a 2’3’-cyclic phosphate and the 3’ exon with a 5’-hydroxyl group (13,15) (Figure 1). TSEN15 and TSEN54 are non-catalytic structural proteins but their precise role in splicing is unknown (13,16). Following cleavage by the TSEN complex, the tRNA exons are joined together by ligation using two distinct pathways, a healing and sealing pathway in plants and *S. cerevisiae* (17) and a direct ligation pathway in metazoans that utilizes the RNA ligase RTCB (10,18,19).

**Figure 1.**
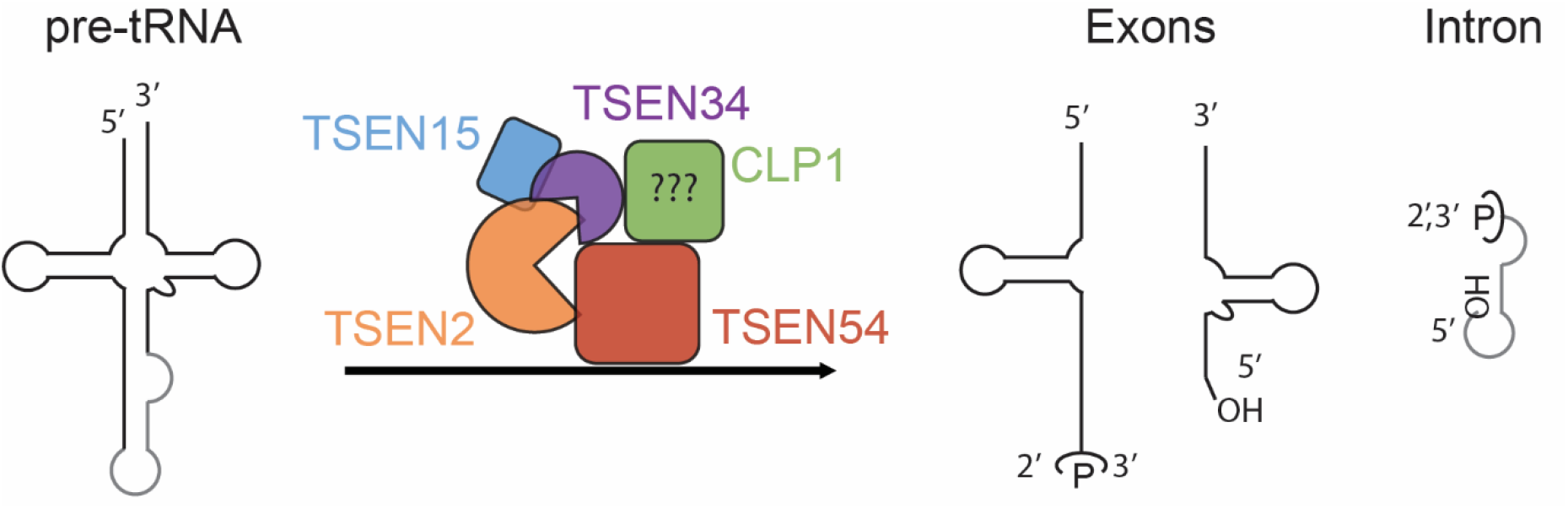
Cleavage of pre-tRNA by the TSEN Complex, the current model of tRNA cleavage. The human TSEN complex is composed of four subunits including two endonucleases (TSEN2 and TSEN34) and two structural components (TSEN15 and TSEN54). Current models for tRNA splicing in metazoans suggest that the TSEN complex requires assistance from CLP1 to catalyze tRNA intron removal (light gray). Following cleavage by the TSEN complex, the 5ꞌ exon and intron contain a 2ꞌ,3ꞌ - cyclic phosphate and the 3ꞌ exon and intron contain a 5ꞌ-OH.

Within the human genome there are 32 predicted intron-containing tRNAs (9). Recent advances in tRNA sequencing confirmed the expression of 26 of the 32 tRNA genes in HEK293 cells (20). The human introns are all found within the same position of the pre-tRNA, one base following the anticodon, between nucleotides 37 and 38 of the mature tRNAs (Figure S1). Eukaryotic pre-tRNAs have a distinctive base pair, known as the proximal base pair, that is absolutely critical for cleavage at the 3’ splice site (Figure S1) (21,22). The eukaryotic TSEN complex has also been proposed to utilize a “ruler” mechanism whereby TSEN54 recognizes features in the mature tRNA domains are used to recognize the 5’ splice site (10,16).

The TSEN complex is vital for eukaryotic life (19,23). Recent work in *S. cerevisiae* revealed that each individual TSEN subunit is essential, even when all intron-containing tRNAs are replaced with intronless variants, suggesting that the complex has a fundamental cellular role beyond tRNA splicing (19). The eukaryotic TSEN complex was first discovered in *S. cerevisiae* in the 1990s (13,24), however it was not until 2004 that the human TSEN complex was identified (14). Intriguingly, upon isolation of the TSEN complex from human cells, an additional TSEN binding partner was identified: the polyribonucleotide 5’-hydroxyl-kinase, CLP1 (14). CLP1 is an established member of the mRNA 3’-polyadenylation machinery (25,26) and is conserved across eukaryotes. The *S. cerevisiae* homologue of CLP1 is an inactive kinase that cannot phosphorylate RNA substrates (27,28), while the CLP1 homologues from *C. elegans* and humans have been shown to be active on RNA substrates *in vitro* (29,30). Mice that express catalytic deficient CLP1 develop a motor neuron disease that can be rescued by transgenic expression of wild-type CLP1 in neurons (31). This finding suggests that CLP1 kinase activity is essential within the nervous system but not in other cell types. The role of CLP1 in tRNA splicing is unclear, but several putative roles for CLP1 have been proposed including maintaining the integrity of the TSEN complex, the existence of a yeast-like ligation pathway, or the promotion of the degradation/turnover of the intron (26).

The human TSEN complex and CLP1 are further linked because of their association with pontocerebellar hypoplasia (PCH), a family of severe neurodegenerative autosomal recessive disorders (23,32–35). There is no curative treatment for PCH and patients rarely survive to adulthood (36). PCH is grouped into 11 subtypes based on phenotypic characteristics such as severe hypoplasia, trouble swallowing, progressive microcephaly, and motor and cognitive impairments (37). A mutation within CLP1 is linked to class PCH10 (34,38,39), while mutations in TSEN2, TSEN15, TSEN34, and TSEN54 are associated with PCH2, PCH4, and PCH5 (32,40). The other PCH subtypes are linked to mutations in other genes, many of which are associated with RNA processing and maturation (37,41–43). The underlying cause of PCH is unknown, but the association of PCH with several RNA processing enzymes suggests a link between disruption of RNA processing and neurological disorders. A homozygous mutation in CLP1 was recently shown to lead to a loss of TSEN complex association and reduced tRNA splicing (34,39). These results led to the hypothesis that, in humans, CLP1 is required for the integrity of the TSEN complex and efficient splicing (26).

The goal of the present study was to reconstitute the core TSEN complex and characterize its activity *in vitro*. Here, we present the full reconstitution of the multi-subunit TSEN complex through use of an *Escherichia coli* co-expression system. The four subunits of the TSEN complex assemble into a stable heterotetramer that actively cleaves intron containing pre-tRNAs as well as a mimic of the pre-tRNA anticodon stem loop. We further show that depletion of the *Drosophila melanogaster* orthologue of CLP1 leads to an increase in production of a mature tRNA reporter construct as well as endogenous and reporter tricRNAs (tRNA intronic circular RNAs). On the basis of these data, we propose that CLP1 is not a critical component of the core TSEN tRNA cleavage machinery but instead functions as a regulatory factor for metazoan tRNA processing.

## MATERIALS AND METHODS

### Recombinant purification of the TSEN Complexes from E. coli

Each individual subunit of the TSEN complex was cloned into a polycistronic vector (pST39)(44). TSEN34, TSEN54, and TSEN2 remained untagged while a C-terminal 6x HIS-tag was cloned onto TSEN15. The TSEN34 (Y247A, H255A, K286A) and TSEN2 (Y369A, H377A, K416A) catalytic mutant co-expression plasmid was generated by Genscript. Please refer to Table S1 for a list of all expression plasmids used in this study. Protein expression for all constructs was conducted using Rosetta II pLac(I) cells (Millipore) grown to high density at 37 °C, cooled to 30 °C, and induced with 2 mM ITPG. Induced cells were incubated for 5-6 hours at 30 °C. Upon the completion of protein expression, cells were harvested by centrifugation and pellets were stored at −80 °C until use.

Cells were re-suspended in Lysis Buffer (50 mM Tris pH 8.0, 10 % glycerol, 500 mM NaCl, 5 mM MgCl_2_, 0.1% Triton-X-100) with the addition of 1 mM phenylmethylsulphonyl fluoride (PMSF). Cells were lysed using sonication and cell lysates were cleared at 15,000 x g for 50 minutes. Clarified lysates were incubated with HIS-60 (Takara) resin for 30-60 minutes, washed with Lysis Buffer, and then eluted in Lysis Buffer plus 250 mM Imidazole. The eluate was injected into a Superdex 200 Increase column (GE) equilibrated with 50 mM Tris pH 8.0, 200 mM NaCl, 5% glycerol, and 5 mM MgCl_2_. Protein was concentrated in a 30 kDa molecular weight cut-off concentrator and then immediately assayed for tRNA cleavage. SDS-PAGE gels were visualized using SimplyBlue SafeStain (Invitrogen).

### Characterization of the TSEN complex using SEC-MALS

Freshly prepared TSEN complex, as described above, was further purified on a Suprose 6 Increase column (GE) equilibrated with 20 mM Tris pH 8.0, 200 mM NaCl, 2% glycerol, and 5 mM MgCl_2_. Protein was then immediately concentrated and analyzed by SEC-MALS as previously reported (45), except using an S200 Increase column (GE) equilibrated with 20 mM Tris pH 8.0, 200 mM NaCl, 2% glycerol, and 5 mM MgCl_2_. An average value and the standard deviation from three independent experiments was calculated.

### Preparation of synthetic RNA substrates

Fluorescently labeled anticodon stem loop RNA was obtained from IDT following HPLC purification (/5Cy5/UUGGACUUCUAGUGACGAAUAGAGCAAUUCAA). Full-length tRNA genes were synthesized and cloned into pUC19 vectors by Genescript, genes were flanked by a 5’ T7 promoter and a 3’ EcoRV restriction site (sequences are available in Table S2). tRNA containing plasmids were linearized using restriction digestion and tRNAs were *in vitro* transcribed overnight using the HiScribe T7 transcription kit (NEB) per the manufacturer’s protocol. RNA was then either cleaned up using the RNAeasy kit (Qiagen) following DNase digestion (Qiagen) or run on TBE-Urea gels (Novex), visualized by UV backlighting on a TLC plate, and purified from the gel. Briefly, the gel was shred through a syringe, RNA was extracted in buffer overnight at 4 °C, gel pieces were removed by filtration through a 0.2 μM filter, and the samples were buffer exchanged into 10 mM Tris pH 8.0 with 1.0 mM EDTA and concentrated. Both methods were suitable for purification of RNA.

### tRNA intron splicing assay

tRNA cleavage reactions were carried out in RNA Cleavage Buffer (50 mM KCl, 50 mM Tris pH 7.5, 5 mM MgCl_2_, 1 mM DTT, and 1 unit/mL RNAsin (Promega)) at room temperature for 30 minutes (unless otherwise noted). Reactions were quenched with 6M Urea-loading dye. Substrates were then stored at 20 °C or boiled at 95 °C for 5 minutes and immediately run on a gel. Samples were separated on 10 % or 15 % TBE-Urea gels (Novex). Gels were then either stained with 1x SYBR Gold (Invitrogen) in 1X TBE, or washed three times in water and stained with 10 μM (5Z)-5-[(3,5-Difluoro-4-hydroxyphenyl)methylene]-3,5-dihydro-2-methyl-3-(2,2,2-trifluoroethyl)-4H-imidazol-4-one (DFHBI-1T)(Tocris) in 50 mM KCl, 50 mM Tris pH 7.5, and 5 mM MgCl_2_. Gels stained by either method were imaged on a Typhoon set to 488 nm.

### tRNA-R1 Northern Blot Protocol

Samples were separated on 15 % TBE-Urea gels (Novex) and then transferred to Amersham Hybond-N+ (GE Healthcare) membrane (equilibrated in 0.5X TBE) using the Trans-Blot Turbo Transfer System (Bio-Rad). Membranes were UV crosslinked with 120 mJ/cm^2^ twice and then pre-hybridized for 2 hrs using DIG Easy Hyb (Roche). Fluorescent end-labeled oligos to the exons of tRNA-R1 were obtained from IDT (See Table S3). The oligo was melted in hybridization buffer at 95 °C and immediately added to the pre-hybridization solution for overnight annealing at 37 °C (5’ exon) or 25 °C (3’ exon) at a final concentration of 1.67 nM. The membrane was then washed 2 x 10 minutes with SSPE Wash 1 (746 mM NaCl, 77 mM NaPO_4_, 6 mM EDTA, 0.1% SDS, pH 7.4) and for 20 minutes with SSPE Wash 2 (77 mM NaCl, 5 mM NaPO_4_, 0.6 mM EDTA, 0.1% SDS, pH 7.4).

### Molecular cloning, expression and isolation of the TSEN complex from HEK cells

Plasmids for pCDNA3.1-GFP-CLP1, pCDNA3.1-TSEN2-FLAG, pCDNA3.1-GFP-TSEN34, and pCDNA3.1-TSEN15-FLAG were obtained from the Genescript EZ orf collection (Ohu26557, Ohu21033, Ohu20820, and Ohu31733, respectively). The DNA for TSEN54 was codon optimized and provided by Genescript. Codon optimized genes for TSEN2, TSEN34, and TSEN15 were also produced by Genescript, including complete active site mutants for TSEN2 and TSEN34. Further subcloning was carried out as follows: TSEN54, and TSEN34 were cloned into pLexM with C-terminal and N-terminal FLAG-tags, respectively. TSEN2 WT and catalytic dead variant (Y369A, H377A, K416A) were also cloned into pLexM with a C-terminal FLAG-tag. TSEN34 and the catalytic dead variant (Y247A, H255A, K286A) were cloned into pCAG-OSF (N-terminal STREP-FLAG tag). Both CLP1 mutants were generated using the NEB Q5 mutagenesis kit and verified by sequencing (Genewiz). A list of all mammalian expression plasmids is available in Table S1.

HEK293 cells were grown in 40 mL suspension cultures, at a density of approximately 2 × 10^6^ cells/mL. Cells were transfected with the aforementioned FLAG or GFP expression plasmids using polyethylenimine (PEI) (2 μg/mL) and grown for approximately 72 hrs, whereupon they were harvested, and frozen at −80 °C until use. Pellets were lysed in HEK IP Buffer (50 mM Tris pH 8.0, 10% glycerol, 100 mM NaCl, 5 mM MgCl_2_, 2 mM EDTA, 0.05% NP-40) with 1 mM PMSF at 4 °C on a nutator for 1 hr. Lysates were cleared by high speed centrifugation at 4 °C. Cleared lysates were then incubated with appropriate resin (Anti-GFP, provided by the NIEHS protein expression core (46)), Anti-FLAG (Pierce™ Anti-DYKDDDDK Magnetic Agarose) and Anti-STREP (MagStrep “type3” XT beads (IBA)), for approximately 60 minutes at 4 °C. Resin was washed 3 times with 1 mL of HEK IP Buffer. If used for RNA cleavage reactions, samples were equilibrated into RNA Reaction buffer immediately prior to performing assays.

### Western blots

Western blots were performed using standard procedures. Protein samples were separated on 4–20% Mini-PROTEAN ® TGX™ precast gels (Bio-Rad) at 300 V for 18-20 minutes and transferred to 0.2 μM PDVF (Bio-Rad) using the Trans-Blot Turbo Transfer System (Bio-Rad). Odyssey blocking buffer (TBS) and fluorescent secondary antibodies (all used at 1:5,000) were purchased from Licor. Blots were developed using a 2-minute exposure on the Licor FC, unless otherwise noted. The following conditions were used for primary antibodies: FLAG (SIGMA-F7425): 1:3,000, GFP (ROCHE-11814460001): 1:3,000, β-Actin (Abcam-ab8224): 1:20,000.

### Drosophila cell culture, RNAi, and in-gel staining

S2 cells were maintained in SF-900 serum-free medium (Gibco) supplemented with 1% penicillin-streptomycin and filter sterilized. S2 RNAi was performed as previously described (47) for 10 days, with dsRNA targeting Gaussia luciferase used as a negative control. The Broccoli dual reporter (22) was transfected on day 7, and cells were harvested on day 10. The reporter (2.5 μg plasmid DNA per well) was transiently transfected using Cellfectin II transfection reagent (Invitrogen) according to the manufacturer’s protocol. Primers used to make PCR products for *in vitro* transcription (to make dsRNA) can be found in Table S4. RNA was isolated using TRIzol Reagent (Invitrogen), with a second chloroform extraction and ethanol rather than isopropanol precipitation (48). To test knockdown efficiency, total RNA was treated with TURBO DNase (Invitrogen) and then converted to cDNA using the SuperScript III kit (Invitrogen) with random hexamer priming. Primers for cbc and 5S rRNA can be found in Table S5.

To visualize Broccoli, RNA samples (5 μg) were electrophoresed through 10% TBE-Urea gels (Invitrogen). Gels were washed three times with dH_2_O to remove Urea and then incubated in DFHBI-1T staining solution (40 mM HEPES pH 7.4, 100 mM KCl, 1 mM MgCl_2_, 10 μM DFHBI-1T (Lucerna)). Following staining, gels were imaged on an Amersham Typhoon 5. To visualize total RNA, gels were washed three times in dH_2_O, stained with ethidium bromide, and imaged on an Amersham Imager 600.

### Northern blotting of Drosophila RNA

RNA samples (5 μg) were electrophoresed through 10% TBE-URA gels (Invitrogen). Following electrophoresis, RNA was transferred to a nylon membrane (PerkinElmer). The membrane was dried overnight and UV-crosslinked. Pre-hybridization was carried out in Rapid-hyb Buffer (GE Healthcare) at 42 °C. Probes were generated by end-labeling oligonucleotides (IDT) with γ-^32^P ATP (PerkinElmer) using T4 PNK (NEB), and then probes were purified using Illustra Microspin G50 columns (GE Healthcare). Upon purification, probes were boiled, cooled on ice, and added to the Rapid-hyb buffer for hybridization. After hybridization, the membrane was washed in saline-sodium citrate (SSC) buffer. For probe sequences, see table S3. Washing conditions are as follows. U1: hybridization at 65 °C, washes (twice in 2x SSC, twice in 0.33x SSC) at 60 °C. Dual reporter probe and tric31905 probes: hybridization at 42 °C, two washes in 5x SSC at 25 °C, two washes in 1x SSC at 42 °C. For the dual reporter probe, two additional washes in 0.1x SSC at 45 °C were performed. After washes, the membrane was exposed to a storage phosphor screen (GE Healthcare) and imaged on an Amersham Typhoon 5.

## RESULTS

### Reconstitution of the human TSEN complex

We have developed a reconstitution protocol for the purification of the human TSEN complex. The active eukaryotic TSEN complex has been successfully isolated from yeast, *Xenopus*, and mammalian cells (13,14,49). However, purification of the TSEN complex from native sources leads to heterogenous mixtures of the core components and variable amounts of known associating factors, such as CLP1 and components of the Pol II 3’ cleavage and polyadenylation machinery (14). In order to define the components necessary for intron cleavage, we developed a reconstitution system for the core TSEN machinery (TSEN2, TSEN15, TSEN34, and TSEN54). Our first approach was to purify the individual TSEN components by expressing them in *E. coli*; however only TSEN15 was found to be soluble on its own. We hypothesized that the other TSEN components may depend upon one another for protein stability. Therefore, we engineered a polycistronic vector for the simultaneous expression of all four TSEN proteins (Figure 2A). We fused a hexa-histidine tag onto the C-terminus of TSEN15 so that the entire complex could be isolated with a nickel affinity column. The eluate from the affinity column was then run over a size exclusion column.

**Figure 2.**
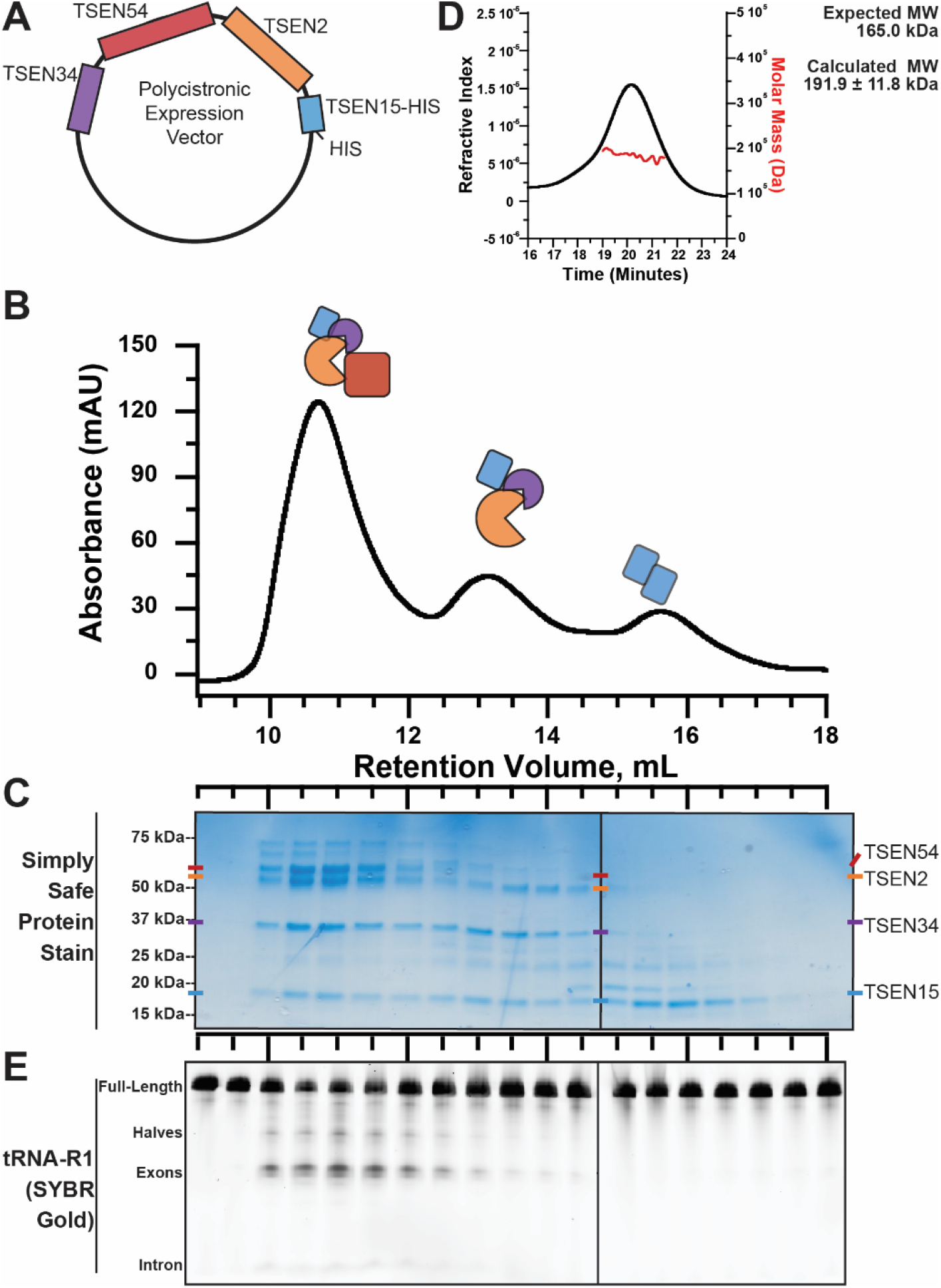
The heterotetrameric human TSEN complex retains nuclease activity in the absence of CLP1. A) Expression of the human TSEN complex in *E*.Coli was possible through use of a polycistronic vector in which each TSEN protein is under independent ribosome binding sites. B) Separation of the TSEN complex via gel filtration produces three distinct TSEN peaks. C) A protein gel reveals the peaks contain the heterotetramer, a trimer (TSEN2, TSEN34, and TSEN15), and pool of soluble TSEN15. D) Cleavage activity assays were performed on fractions from gel filtration separation of the TSEN complex. Cleavage reactions were separated on denaturing TBE-Urea gels, which were stained with SYBR Gold to visualize total RNA. E) SEC-MAL analysis of the peak centered around 10.5 mL reveals the TSEN complex is a single heterotetramer with an average mass of 191.9 kDa.

We observed three prominent peaks from our size-exclusion profile located at retention volumes of 10.5 mL, 13.5 mL, and 15.5 mL (Figure 2B). We analyzed these peaks by SDS-PAGE to determine their protein composition (Figure 2C). The first peak at 10.5 mL contains all four subunits (TSEN2, TSEN15, TSEN34 and TSEN54). The identities of TSEN2, TSEN15, TSEN34, and TSEN54 were confirmed by mass spectrometry and the SDS-PAGE analysis is suggestive of an equimolar ratio of each subunit. To confirm the oligomeric state of the core complex, we analyzed the 10.5 mL peak fraction by size-exclusion chromatography, paired with in-line multi-angle light scattering (SEC-MALs) (Figure 2D). We measured an average molecular weight of 191.9 ± 11.8 kDa from three independent purifications, all with a polydispersity value of 1.0. These findings are consistent with the theoretical molecular weight of the TSEN heterotetramer. The second peak at 13.5 mL contains TSEN2, TSEN15, and TSEN34, suggesting that these components can form a stable sub-complex in the absence of TSEN54. The final peak at 15.5 mL contains TSEN15 alone. Previous work with the isolated TSEN15 subunit revealed that it forms a homodimer on its own (50), thus we infer that this peak represents a TSEN15 homodimer.

Next, we confirmed that the recombinant TSEN complex retained cleavage activity. To verify tRNA splicing endonuclease activity we transcribed the human tRNA-R1 (tRNA^ARG-TCT)^ transcript *in vitro*, which includes a 15-nucleotide intron (Figure S1). Expression of this gene was recently validated in human cells using a new tRNAseq methodology (20). The pre-tRNA was incubated with individual fractions from the SEC column and the cleavage reactions were analyzed on denaturing TBE-Urea gels. Gels were stained with SYBR Gold and visualized with a fluorescent imager. Prominent tRNA cleavage products were visible with the fractions corresponding to the full heterotetrametric TSEN complex but not for any of the other fractions (Figure 2E). This result confirms that our *E. coli* co-expression system produces the biochemically active TSEN complex and that, similar to studies of the *S. cerevisiae* TSEN complex (13), the human heterotetrametric TSEN core is sufficient for intron cleavage.

### Human TSEN core is active on pre-tRNA substrates

We characterized the endonuclease cleavage activity of the recombinant TSEN complex and found that it is active on multiple intron-containing tRNA substrates. First, we carried out a time course with full-length *in vitro* transcribed pre-tRNA-R1 (Figure 3A) and the recombinant TSEN complex. Cleavage products were analyzed by denaturing gel and visualized with SYBR Gold (Figure 3A). We confirmed the identity of the cleavage products with Northern blot probes to the 5’ and 3’ ends of tRNA-R1 (Figure S2). To ensure that the cleavage products were generated by the TSEN complex and not a contaminant from our *in vitro* system, we created a catalytic deficient AAA^2^-TSEN variant. Sequence alignments across TSEN endonucleases suggests that the endoribonuclease active sites are well conserved (Figure S3A) (15,51,52). Each endoribonuclease active site is composed of three invariant residues including a histidine, lysine, and tyrosine. A structure of the homodimeric tRNA splicing endonuclease machinery from *Archaeoglobus fulgidus* bound to RNA revealed that these invariant residues cluster around the cleavage site and suggests that they support catalysis through a general acid-base mechanism (53). To inactivate the human TSEN endonuclease subunits, TSEN34 (3’ site) and TSEN2 (5’ site), we made triple alanine substitutions of the catalytic triad from each active site; TSEN34 (Y247A, H255A, K286A) and TSEN2 (Y369A, H377A, K416A). The AAA^2-^TSEN complex could be purified suggesting that the active site mutants do not interfere with the assembly of the TSEN complex (Figure S3B). We performed the tRNA-R1 time course with AAA^2-^TSEN and did not observe cleavage products by either SYBR Gold staining (Figure 3A) or Northern blot (Figure S2B). Collectively, these results confirm that the specific cleavage products observed in our assays are from the active TSEN complex.

**Figure 3.**
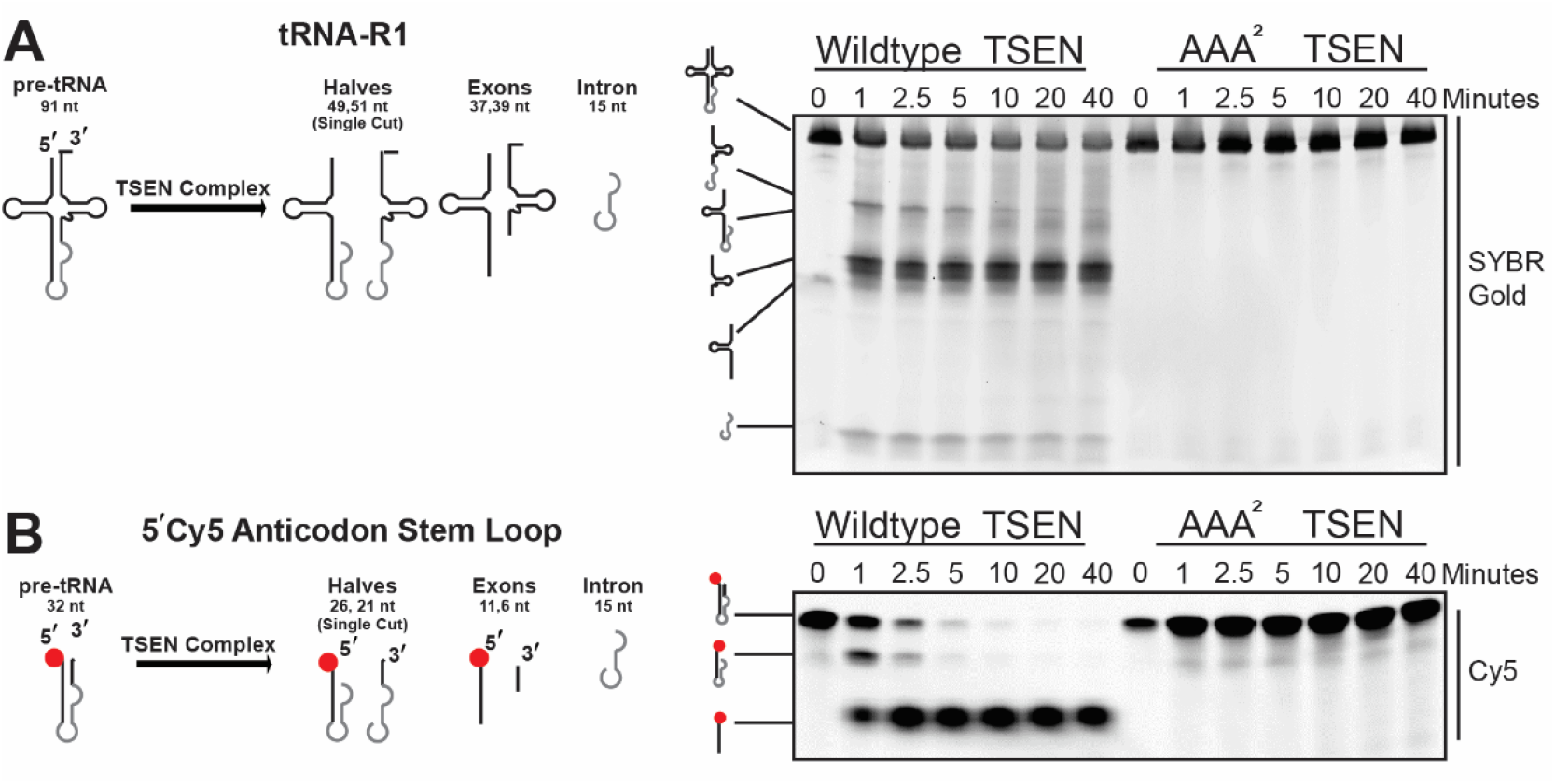
Cleavage of tRNA substrates by the human TSEN complex. The heterotetrameric TSEN complex is efficient at cleaving A) tRNA-R1 full-length (1.7 μM) and B) anticodon stem loop (0.5 μM) tRNA substrates. Representative time courses (0, 1, 2.5, 5, 10, 20, 40 minutes) are shown for the cleavage of each tRNA substrates by 1.5 μM TSEN complex (both wild-type and endonuclease dead (AAA^2^) complexes). RNA cleavage products were separated on TBE-Urea denaturing gels and visualized using a typhoon.

After confirming activity on a full length pre-tRNA, we found that the recombinant TSEN complex is also able to cleave a truncated pre-tRNA substrate. Due to a lack of structural information for the eukaryotic TSEN complex, our understanding of substrate recognition by the complex is limited. Previous work suggests that the eukaryotic TSEN complex has broad specificity. For example, the Xenopus TSEN complex can cleave a bulge-helix-bulge (BHB) substrate, the universal substrate for archaeal TSENs (49), whereas recent work with the *S. Cerevisiae* TSEN complex identified the stem loop of CBP1 mRNA as a new TSEN substrate (54). To begin to address substrate recognition, we designed a synthetic mini intron construct with a 5’ fluorescent label (Figure 3B), that includes only the anticodon stem loop (ASL) of tRNA-R1 (Figure S1). Using this substrate, we can detect the uncleaved product, the 5’ half, and the 5’ exon (Figure 3B). We carried out a time course of the cleavage reaction with the ASL substrate and observed robust and efficient cleavage with the WT-TSEN complex, but not with the catalytic site mutant (Figure 3B). Therefore, the human TSEN complex does not require the full tRNA cloverleaf structure to efficiently process the anticodon stem *in vitro*.

Finally, we also characterized the activity of the recombinant TSEN complex using an alternative pre-tRNA substrate. Due to its small size, the excised wild-type tRNA intron is difficult to detect by conventional methods. Therefore, we designed a synthetic tRNA-I21 (tRNA^ILE-TAT^) variant bearing a Broccoli-tagged intron (tRNA-I21-broccoli) (55,56) (Figure S4). The Broccoli aptamer binds to a fluorescent dye called DFHBI-1T, such that only intron(broccoli)-containing RNAs are visualized (55). Thus, in DFHBI-1T stained gels, only pre-tRNA, intron-containing tRNA halves, and introns are detected (Figure S4). We carried out a time course of the cleavage reaction with tRNA-I21-broccoli with the recombinant TSEN complex. Cleavage products were resolved by denaturing gel electrophoresis, refolded in the gel, stained with DFHBI-1T, and visualized with a fluorescent scanner. Through this approach, the intron cleavage product was clearly detectable with the wild-type TSEN complex (Figure S4). Collectively, these results establish that our recombinant TSEN complex is active on a variety of intron containing pre-tRNA substrates *in vitro*.

### CLP1 is not required for tRNA cleavage

To address questions regarding the role played by the polynucleotide kinase, CLP1, in the TSEN complex, we first attempted to form the CLP1•TSEN complex using a recombinant *E. coli* system. However, we could not generate enough of the CLP1•TSEN complex for biochemical experiments. This result suggests that there may be additional factors or post-translational modifications that mediate CLP1•TSEN association. To purify the CLP1•TSEN complex, we developed a large-scale mammalian expression system to isolate the TSEN complex bound to CLP1. To first establish the recombinant mammalian system, we individually and co-expressed each TSEN protein harboring a FLAG tag and then carried out immunoprecipitations using anti-FLAG resin (Figure 4A). Activity assays on our resin-bound isolates revealed detectable tRNA cleavage only following co-expression of all four TSEN subunits (Figure 4B). Correspondingly, we were unable to detect the presence of stoichiometric amounts of the individual endogenous TSEN proteins by SDS-PAGE analysis of isolates stained with SYPRO Orange (Figure 4C). The lack of nucleolytic activity as well as the absence of the full tetrameric TSEN complex within these immunoprecipitates suggests that, as in our recombinant *E. coli* system, co-expression of all four subunits is required to stably isolate the full TSEN complex cultured in human cells.

**Figure 4.**
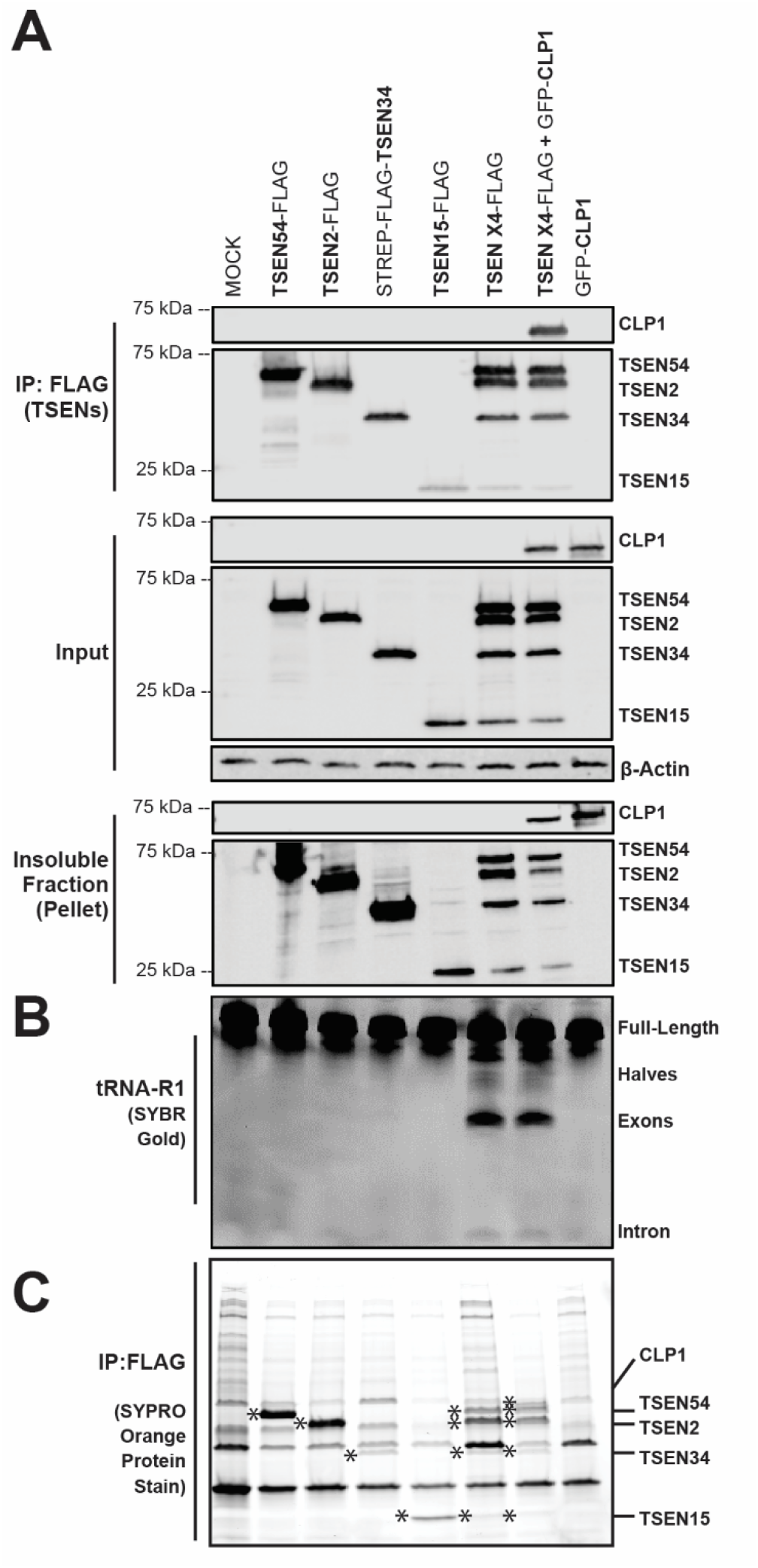
Co-expression and Immunoprecipitation of the TSEN complex from HEK cells yields an active complex. A) FLAG-tagged TSEN constructs were transiently co-expressed and immunoprecipitated, using Anti-FLAG resin, from HEK 293 cells. Isolated samples (IP), soluble cell lysates (input), and the insoluble fraction (pellet) were analyzed by Western blot. TSEN subunits were detected with an anti-FLAG antibody, CLP1 was detected with an anti-GFP antibody, and β-actin was used as a loading control. B) Denaturing TBE-Urea gels were used to analyze immunoprecipitated samples assayed for endonuclease activity on pre-tRNA-R1, the gel was stained with SYBR Gold. C) Finally, IP samples were separated on an SDS-PAGE gel and stained to visualize proteins using SYPRO Orange protein stain. TSEN proteins are denoted by “*” in respective lanes.

To ensure that the observed cleavage was specifically from our overexpressed complexes and not endogenous TSEN subunits, we assayed tRNA cleavage using the recombinant TSEN complex with either wild-type or triple alanine endonuclease-dead variants of TSEN34 (Y247A, H255A, K286A) and TSEN2 (Y369A, H377A, K416A). Cells were transfected with plasmids harboring FLAG-tagged TSEN2, TSEN15, and TSEN54 along with Strep-FLAG tagged TSEN34. The active tetrameric TSEN complex was successfully isolated by co-immunoprecipitation with Strep-FLAG-TSEN34 and we observed no non-specific binding of the other subunits to the anti-Strep resin (Figure S5A). Moreover, mutation of the active sites within TSEN34 and TSEN2 did not impair TSEN protein expression or TSEN complex formation (Figure S5A, top two panels). We assayed the Strep-bound isolates with pre-tRNA-R1 and detected cleavage products by denaturing gel. We observed accumulation of the 3’ exon half with the TSEN34-AAA mutant, whereas accumulation of the 5’ exon half was observed with the TSEN2-AAA mutant (Figure S5B). Co-expression of both active site mutants led to a loss of observable cleavage products (Figure S5B).

After establishing our HEK293 recombinant system, we formed the TSEN-CLP1 by co-transfecting cells with plasmids harboring all four TSEN subunits and CLP1. Using this approach, we could isolate the TSEN•CLP1 complex by co-immunoprecipitation with FLAG-tagged TSEN subunits (Figure 4A) or Strep-tagged TSEN34 (Figure S5). We saw no observable improvement in tRNA cleavage when GFP-CLP1 was also overexpressed with the TSEN complex (Figure 4A, lane 6 vs 7; Figure S5, lanes 1-6 versus lanes 7-12), suggesting that CLP1 does not play a central role in regulating tRNA cleavage within the mammalian TSEN complex *in vitro*.

### The *Drosophila* CLP1 orthologue, cbc, is a negative regulator of tRNA exon ligation and intron circularization *in vivo*

Given our finding that CLP1 is not necessary for TSEN-mediated tRNA intron removal *in vitro*, we sought to determine whether CLP1 might play a downstream role in tRNA processing. Previous work has shown that the CLP1•TSEN complex can cleave pre-tRNAs and phosphorylate both the 3’ tRNA exon and the intron (29). Phosphorylation of the exon prevents RTCB mediated ligation of the two exon halves, whereas phosphorylation of the intron prevents intron circularization (18). We harnessed an *in vivo* system to evaluate the role of CLP1 in tRNA splicing. We previously identified the *Drosophila melanogaster* TSEN orthologs using a cell culture-based assay in which a tRNA:Tyr reporter construct containing a Broccoli aptamer in its intron was assayed for both intron circularization and tRNA maturation (22). We used this ‘dual reporter’ to determine if CLP1 activity is important for tRNA intron removal and maturation in metazoan cells. We carried out RNA interference against the *Drosophila* orthologue of CLP1(called crowded by cid, or cbc) in S2 cells, and verified cbc knockdown by RT-PCR (Figure 5A). We then measured the efficiency of tRNA intron removal by examining the formation of tricRNAs, which are a known *in vivo* product of the tRNA splicing pathway (22,57). We observed a striking increase in levels of both the endogenous tric31905 RNA (Figure 5B) as well as the tricBroc reporter (Figure 5C). These data suggest that CLP1/cbc also plays an inhibitory role in tricRNA biogenesis *in vivo*.

**Figure 5.**
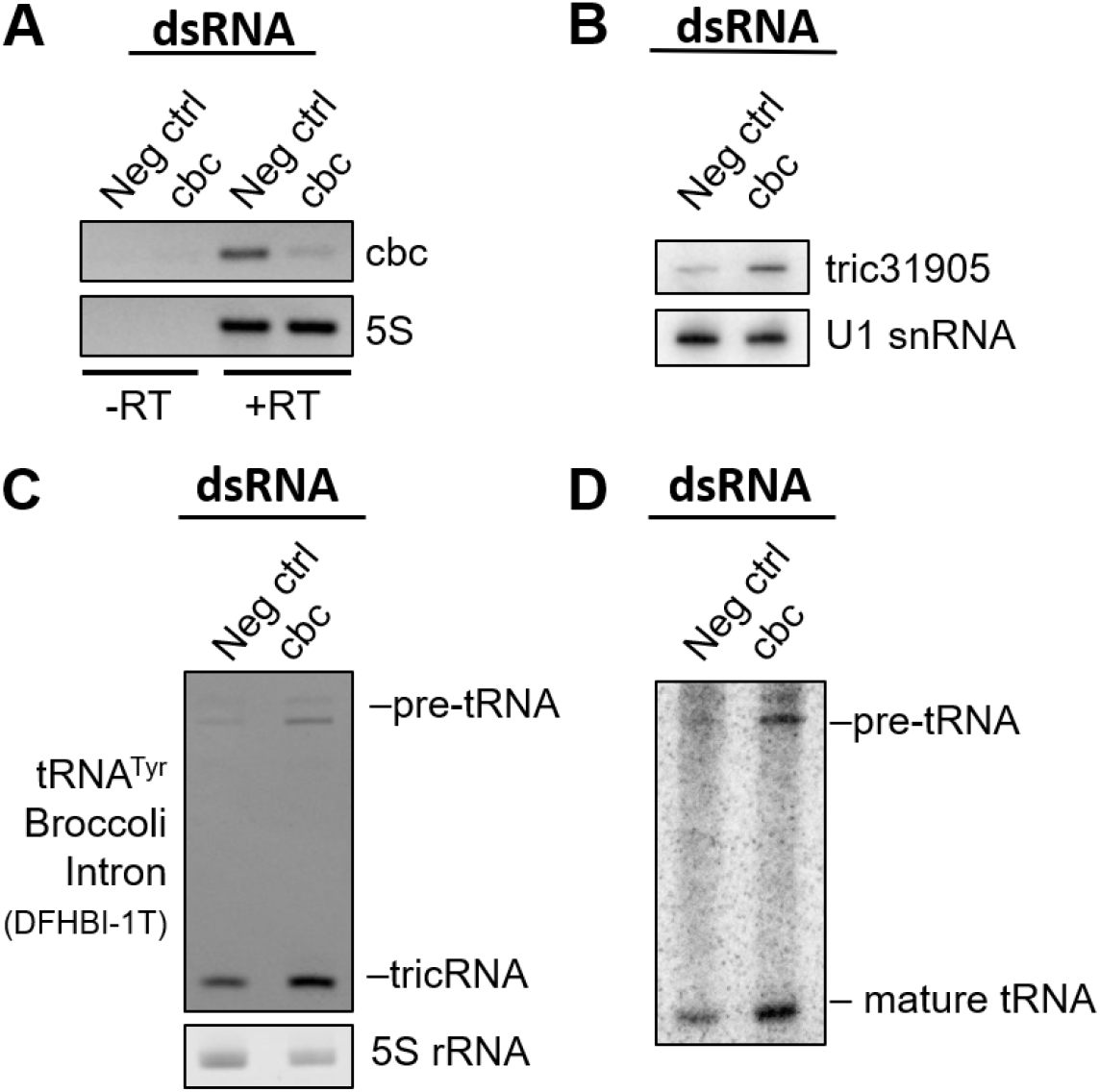
Depletion of the *Drosophila melanogaster* CLP1 ortholog, cbc, in S2 cells does not prevent the generation of tricRNAs or tRNA maturation. S2 cells were treated with dsRNA to knock down cbc *in vivo*. Cells were then transfected with a *D. melanogaster* tRNA-Tyr reporter containing Broccoli in the intron (72). Total RNA was isolated from the cells. A) RT-PCR was used to measure successful cbc knockdown by dsRNAs, with 5S rRNA and unamplified RNA as controls. B) A northern blot probing tric31905 was performed to measure levels of endogenous circular intron production. Levels of the snRNA U1 were used as a control for the northern blot. C) RNA was run on a TBE-Urea gel and visualized using DFHBI-1T. The gel was then re-stained with ethidium bromide, and the 5S rRNA is shown as a loading control. D) A northern blot probing the reporter tRNA was performed to measure levels of tRNA maturation. The U1 snRNA control from panel B is the same control for this northern blot.

Previously, we showed that RTCB is the ligase responsible for tRNA intron circularization in *Drosophila* (22,57). Given the wealth of data demonstrating that RTCB-type ligases are inhibited by the presence of a 5’ phosphate (58–61), we hypothesized that CLP1/cbc would also inhibit formation of mature tRNAs. Consistent with this notion, northern blotting for the reporter RNA revealed a similar increase in levels of the mature tRNA:Tyr construct following depletion of CLP1/cbc (Figure 5D). Together with our *in vitro* cleavage data, these *in vivo* results provide strong evidence that CLP1 kinase is neither required for tRNA intron cleavage, nor for exon ligation. We therefore propose that CLP1 and its orthologs are critical negative regulators of tRNA processing in animal cells (Figure 6).

**Figure 6.**
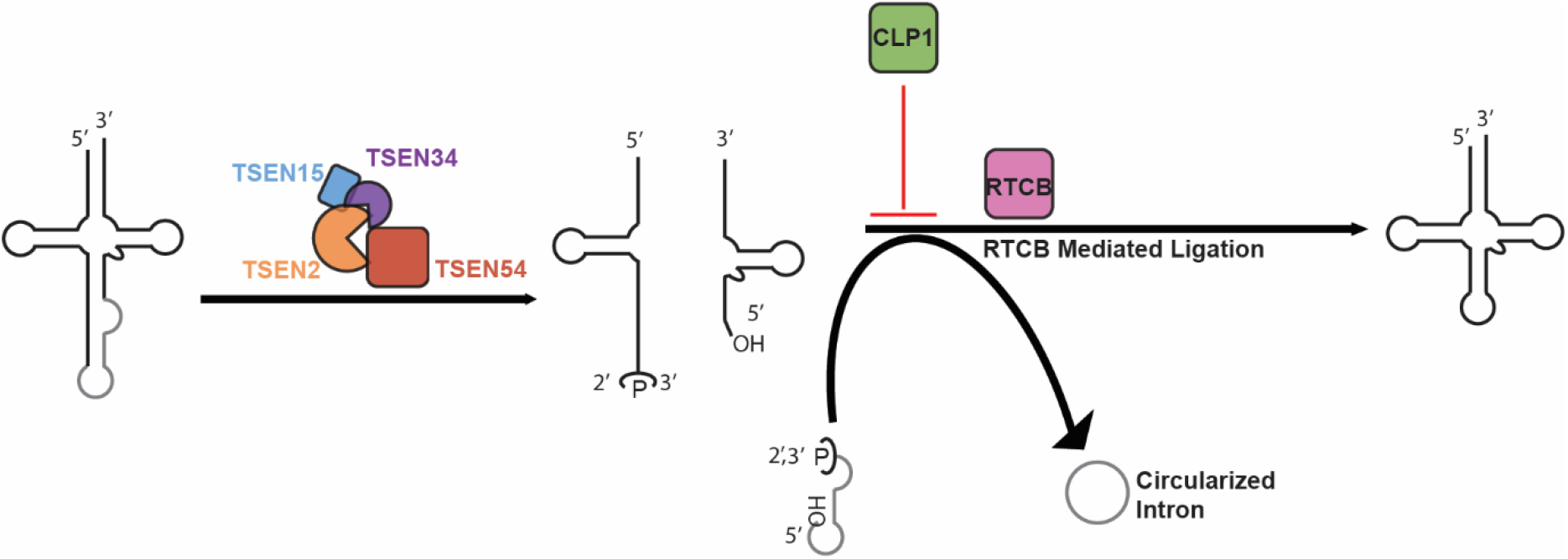
CLP1 is a negative regulator of tRNA splicing. An updated model of tRNA splicing in which the TSEN complex can solely cleave the introns from pre-tRNAs and CLP1 plays a primary role in regulating the RTCB mediated ligation step of splicing.

## DISCUSSION

Our data provide a rational for the formation of the CLP1•TSEN complex in metazoans. Prior to this work, several putative roles for CLP1 in tRNA splicing have been proposed, including: supporting the integrity of the TSEN complex, the existence of an alternative yeast-like ligation pathway, turnover of the intron, or processing of non-tRNA substrates (26,27,29,34,39). Our reconstitution system demonstrates that the heterotetrametric TSEN core is functional in the absence of CLP1. Moreover, purification of the TSEN complex from transfected human cells in the presence or absence of CLP1 does not cause observable changes in tRNA splicing. This finding suggests that the core tRNA processing machinery is conserved from yeast to humans (13). Unity amongst the eukaryotic TSEN complexes is further supported by our confirmation of the TSEN2 and TSEN34 active sites and the stoichiometry of the complex as a heterotetramer. While we cannot rule out putative roles of CLP1 in tissue-specific and/or spatial regulation of the TSEN complex *in vivo*, our work supports the hypothesis that eukaryotic TSEN complexes catalyze the excision of tRNA introns through a conserved mechanism.

In contrast to the unity we show for the eukaryotic splicing machinery, ligation of tRNA exons in yeast and humans is thought to be carried out through two distinct pathways. In the budding yeast S. *cerevisiae*, exons are ‘healed and sealed’ by the multicomponent enzyme Trl1, which exhibits cyclic phosphodiesterase, polynucleotide kinase, and RNA ligase activities (11). Trl1 opens the 5’-exon’s cyclic-phosphate termini, phosphorylates the 3’-exon’s 5’-hydroxyl termini, and ligates the ends back together. Both CLP1 and the human phosphodiesterase CNPase were previously shown to be capable of rescuing Trl1 mutants in their respective domains (27,62), but no human tRNA ligase capable of ligating the 3’-exon’s 5’-phosphate to the 3’-hydroxyl has been identified. In contrast, the ‘direct ligation’ pathway in metazoan cells relies on RTCB to ligate the tRNA halves - joining a 5’ hydroxyl directly to the 2’,3’-cyclic-phosphate (31). CLP1-mediated phosphorylation of the 3’-exon’s 5’-hydroxyl would effectively inhibit RTCB-mediated ligation of the exon halves. Thus, CLP1 could either play a role in an alternative “yeast-like” ligation pathway, or, as our data suggest, it could act as a negative regulator of RTCB-mediated ligation, which is believed to be the primary tRNA processing pathway in animal cells (63).

RTCB not only is responsible for tRNA ligation, but it is also responsible for creation of tricRNAs in metazoans and archaea (22,57,64,65). Both metazoans and archaea have been shown to have produce tricRNAs, whereas yeast introns are primarily linear and rapidly degraded (66,67). Because circularized introns are very stable (57), CLP1 may play a critical role in circular RNA homeostasis by inhibiting RTCB-mediated ligation and acting as negative regulator of tricRNA biogenesis. Along these lines, we showed here that knockdown of the *D. melanogaster* CLP1 homolog, cbc, increased the generation of endogenous circularized tRNA introns. Therefore, we propose that CLP1 acts as a critical negative regulator of tricRNA biogenesis and tRNA maturation. Beyond tRNA splicing CLP1 may also function as a negative regulator of other RNA processing pathways dependent upon RtcB ligation such as the unfolded protein response(68–71).

Here, we establish that the human TSEN complex assembles into a heterotetramer that is sufficient to support the cleavage of tRNA introns *in vitro*. We provide data to support the hypothesis that CLP1 is a negative regulator of tricRNA biogenesis, through blocking RTCB-mediated ligation of the excised intron. Mutations in CLP1 and all four subunits of the TSEN complex are associated with the PCH family of severe neurological disorders. However, CLP1 and the TSEN complex associate with distinct PCH subtypes. This suggests that association with PCH development may not just be in tRNA maturation, but also in the processing of other, yet to be characterized, RNAs. The TSEN reconstitution system described here will be useful for future studies and characterization of additional to be discovered RNA targets of the TSEN complex.

## SUPPLEMENTARY DATA

Supplementary Data are available below.

## ACKNOWLEDGEMENT

We thank Drs. Andrew Sikkema and Marcos Morgan as well as members of the Stanley Lab for their critical reading of this manuscript. We would like to acknowledge Robert Petrovich and the NIEHS Protein Expression Facility for assistance with protein expression in HEK293 suspension cultures and for supplying the anti-GFP resin. We also thank Robert Dutcher for his assistance with running SEC-MALS.

## SUPPLEMENTARY DATA

**Figure S1.**
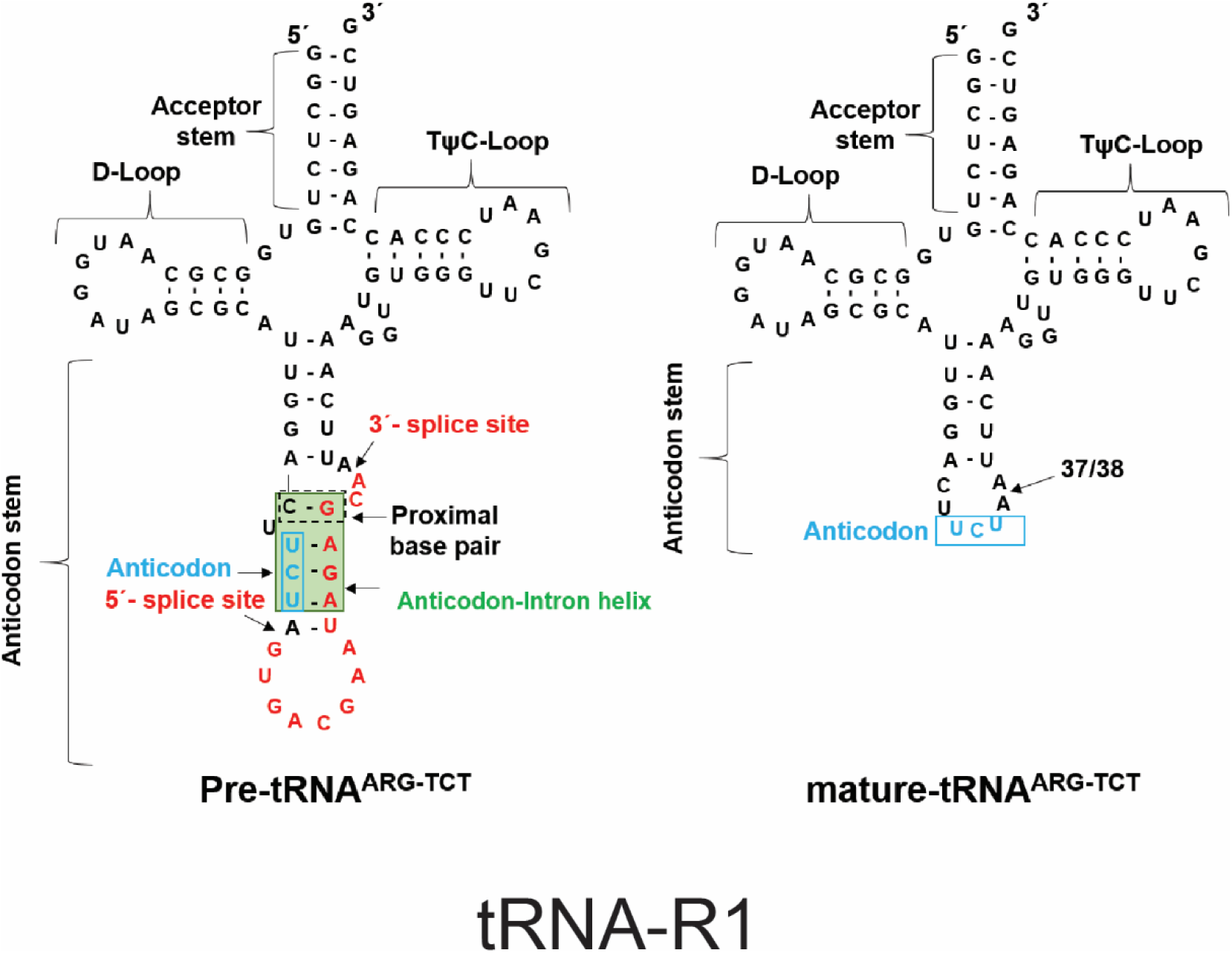
Human tRNA-R1 substrate used in this study. Transfer RNA tRNA-R1 is shown here as pre-tRNA (left) and mature tRNA (right). Key regions of the tRNA are labelled.

**Figure S2.**
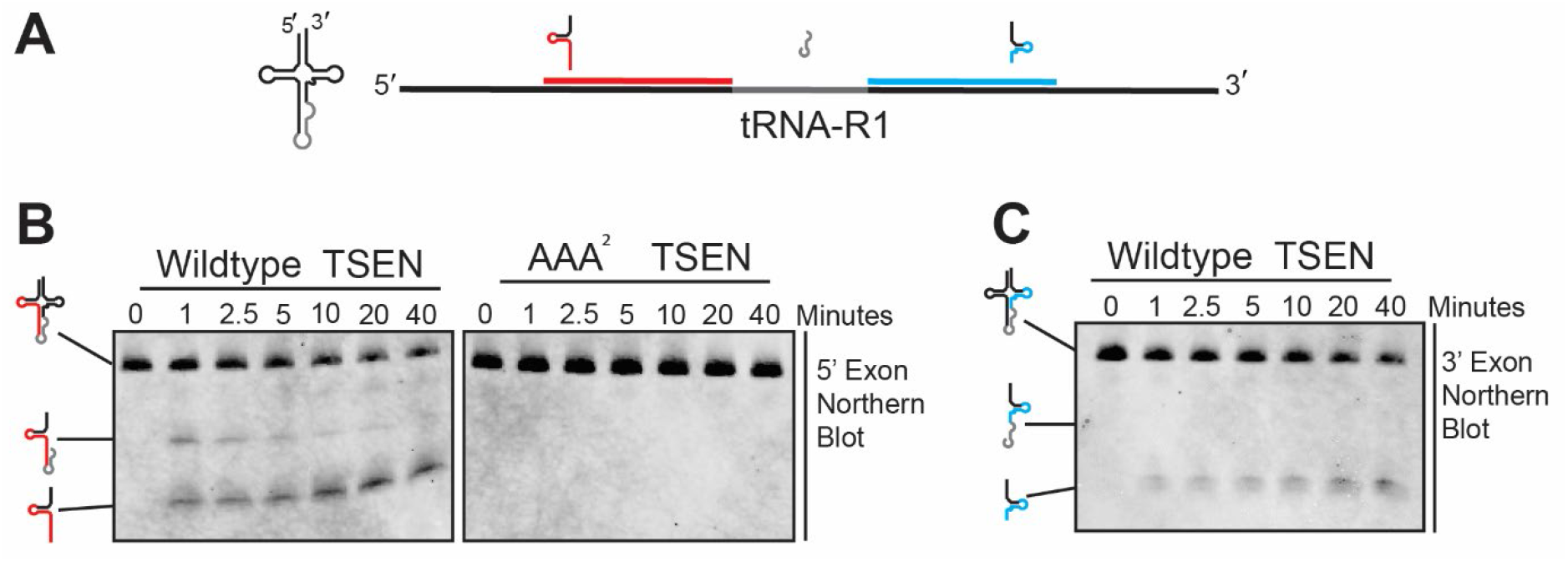
Northern blot validation of time course experiments. Fluorescently labelled DNA oligos specific to tRNA-R1 were designed to bind to the 5’ (red) or 3’ (blue) exons in the regions flanking the intron. A time course of the human TSEN complex (1.5 μM) cleaving tRNA-R1 (1.7 μM) was performed, and samples were analyzed by Northern blotting. Northern blots to the 5’ (B) and 3’ (C) exons were performed.

**Figure S3.**
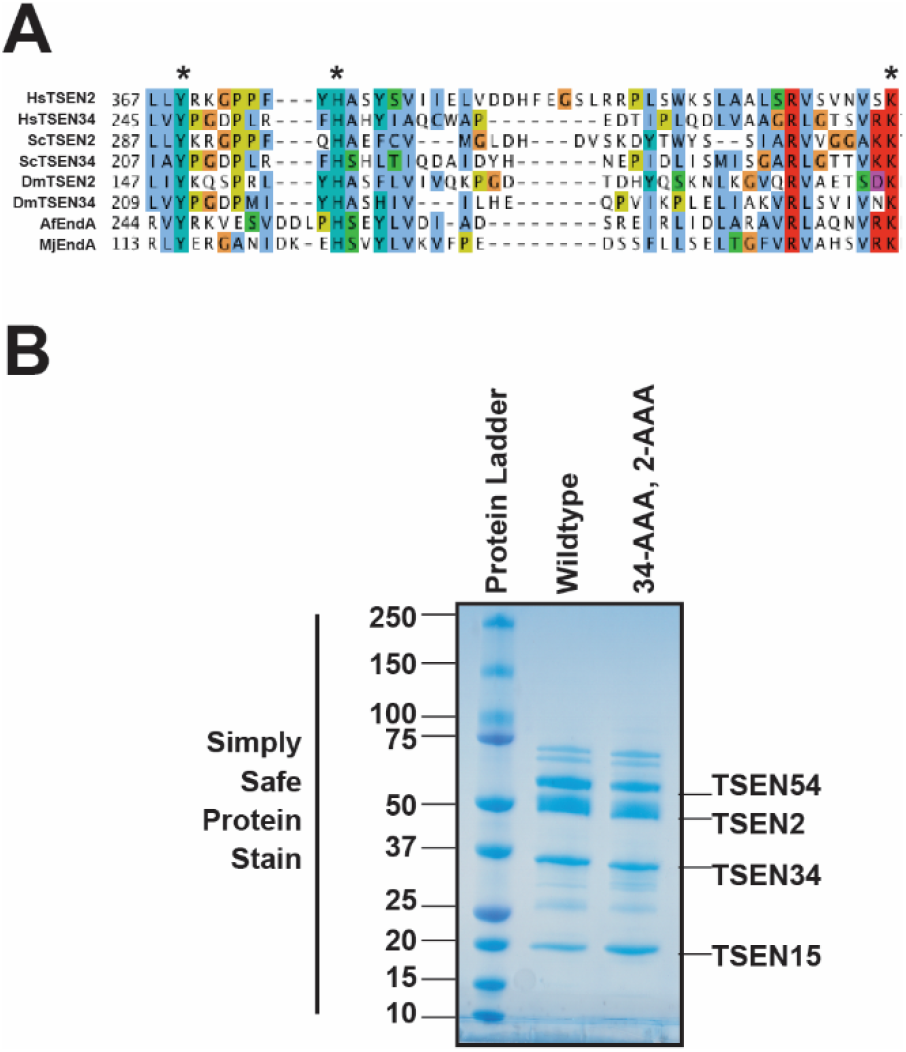
Active site conservation of the TSEN endonucleases. A) A multiple sequence alignment of the active sites for the TSEN2 and TSEN34 endonuclease subunits from *Homo Sapiens (Hs), Saccharomyces Cerevisiae (Sc)*, and *Drosophila Melanogaster (Dm)* shows the conservation of the active site with two archaea (*A.fulgidus (Af) and Methanococcus Jannaschii (Mj))* endonucleases. B) Comparison of 1.875 μg of total protein of Wildtype and AAA^2^ Human TSEN complexes purified from E. coli.

**Figure S4.**
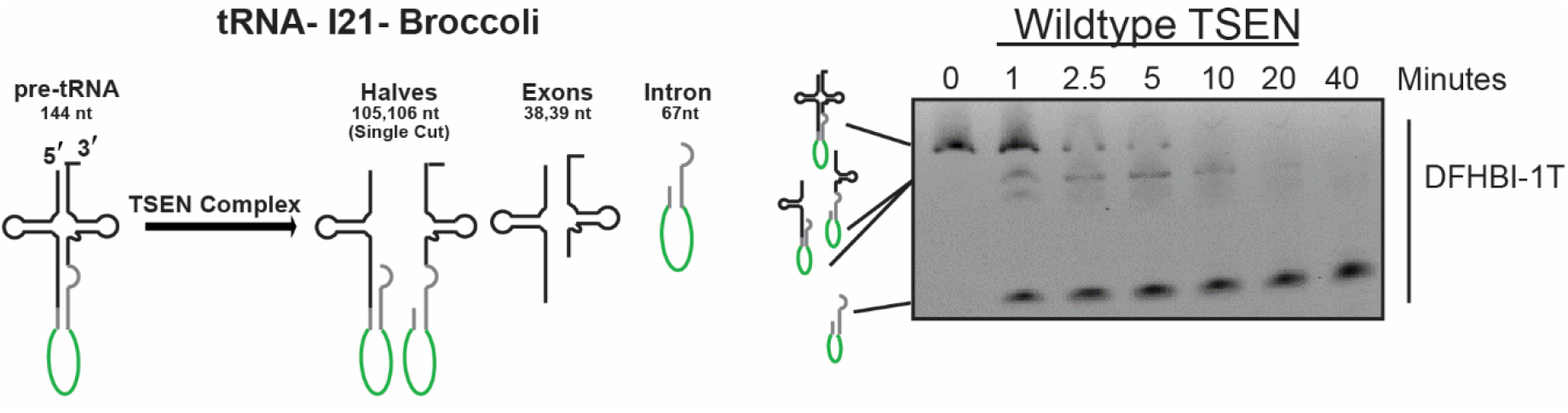
RNA cleavage time course of tRNA substrate tRNA-I21-Broccoli by the human TSEN complex. The *E. coli* recombinant TSEN complex efficiently cleaves the tRNA-I21 with a Broccoli RNA aptamer engineered into its native intron. A time course (0, 1, 2.5, 5, 10, 20, 40 minutes) is shown for the cleavage of the tRNA substrate by 1.5 μM TSEN complex. RNA cleavage products were separated on a TBE-Urea gel, refolded in the gel, stained with 10 μM DFHBI-1T, and -containing RNA visualized using a typhoon imager set to 488 nm.

**Figure S5.**
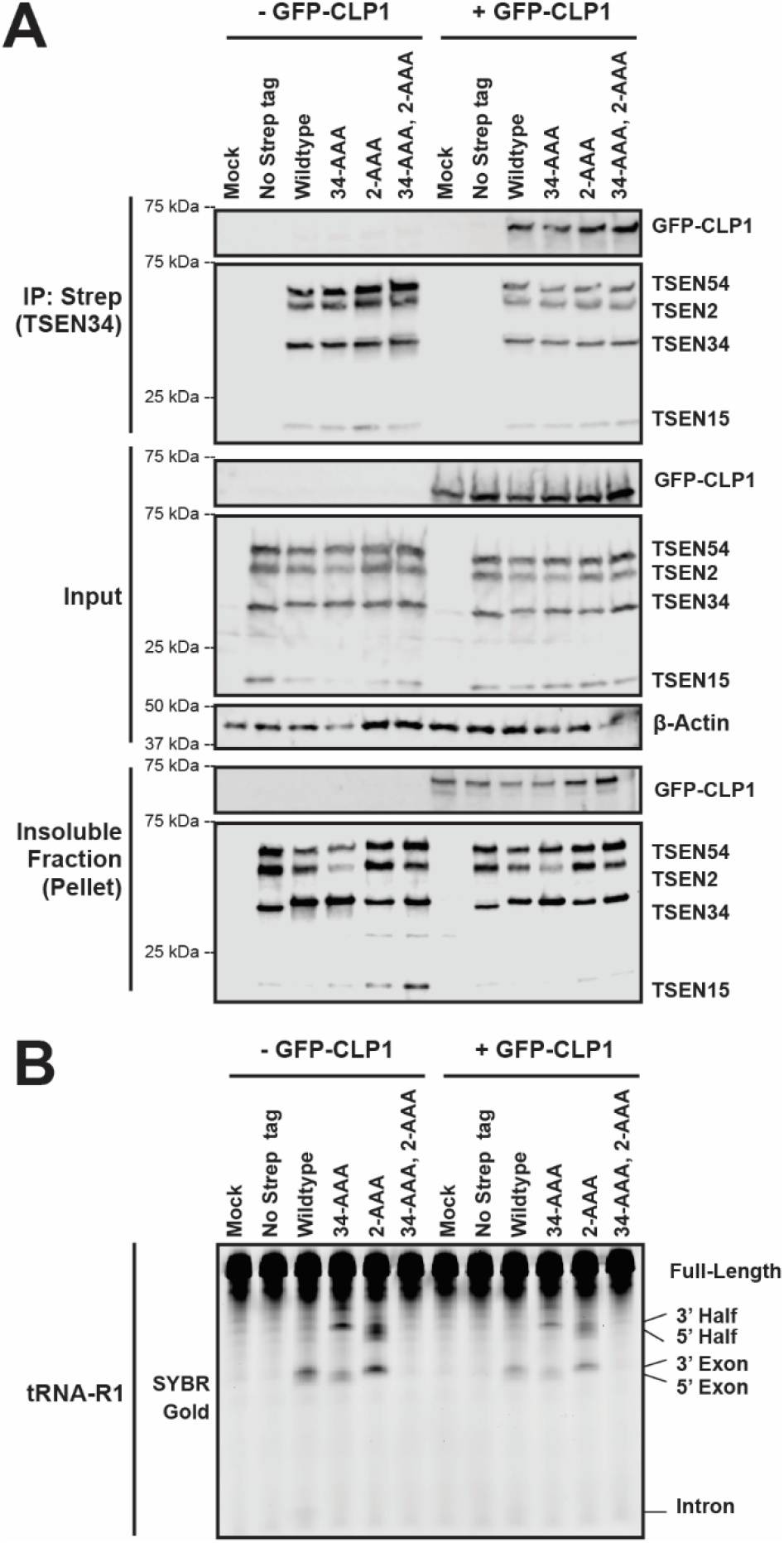
Characterization of tRNA cleavage with TSEN34 and TSEN2 catalytic mutants. A) HEK293 cells were transiently co-transfected with plasmids for FLAG-tagged TSEN subunit variants, including wild-type and endonuclease dead variants of TSEN2 and TSEN34. Samples were co-transfected with or without GFP-CLP1. Samples were immunoprecipitated using Anti-Strep resin to target Strep-FLAG-TSEN34. B) Immunoprecipitated proteins were assayed for cleavage of tRNA-R1. Samples were analyzed by denaturing TBE-Urea gels stained with SYBR Gold to visualize all tRNA products.

**Table S1.**
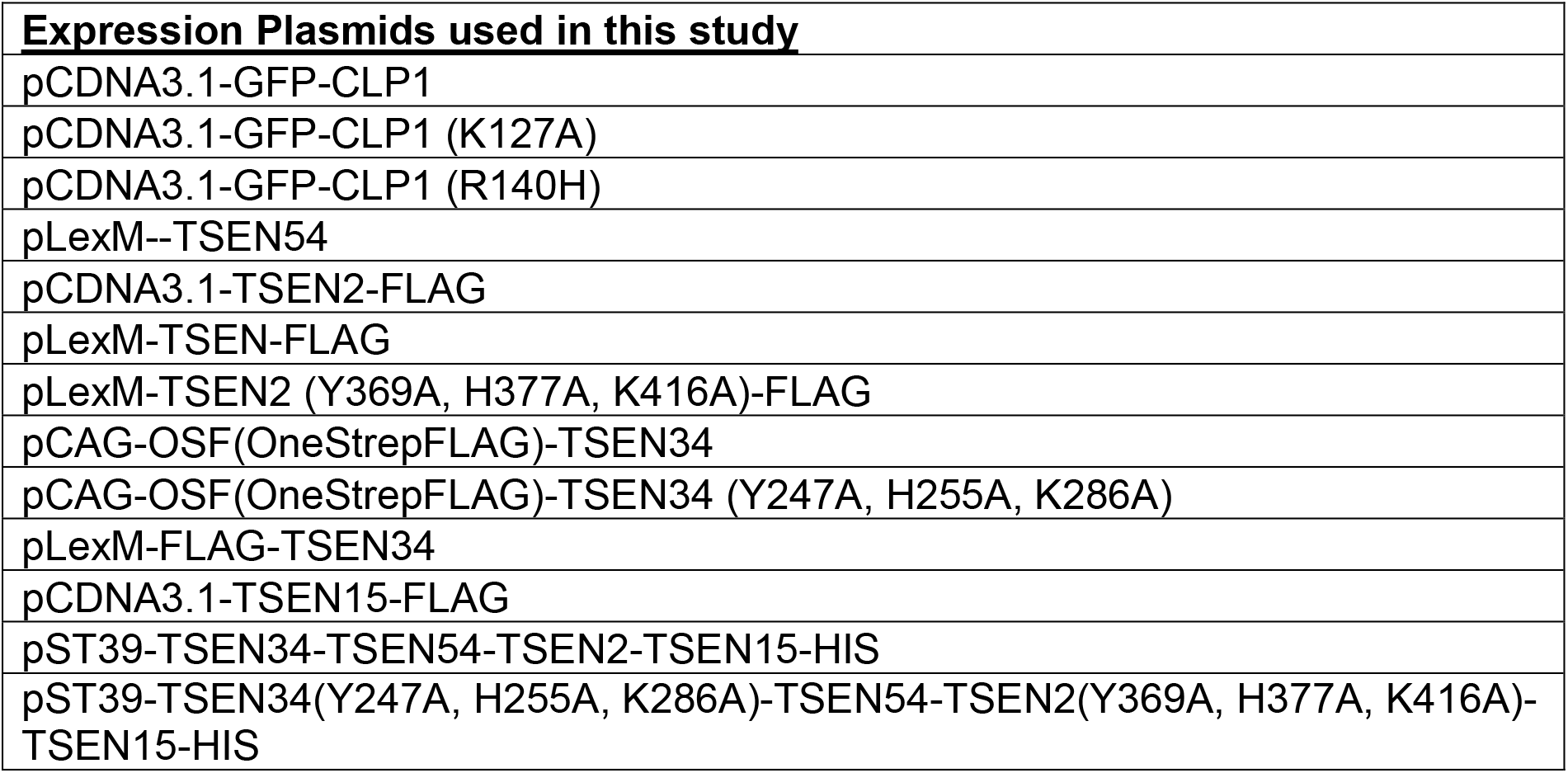
Plasmids used in this study.

**Table S2.**
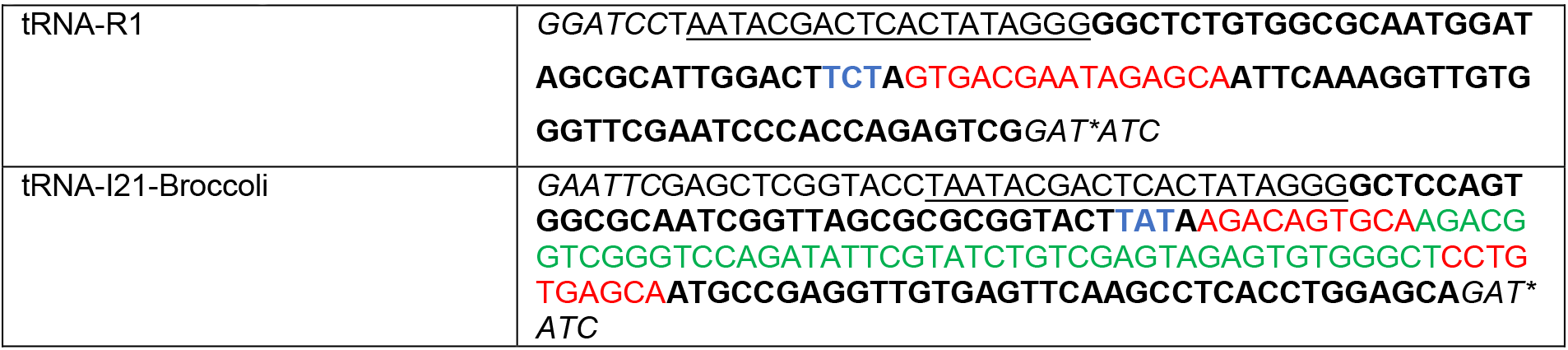
tRNA DNA templates for *in vitro* transcription. Constructs were cloned into pUC19 vectors using BamHI or EcoR1/EcoRV. Vectors are engineered as follows: *BamHI/EcoR1 site*-<u>T7 promoter</u> – **5’ exon**– intron (Broccoli) – **3’ exon**– *EcoRV site (*blunt cut site).* The anticodon is shown in blue.

**Table S3.**
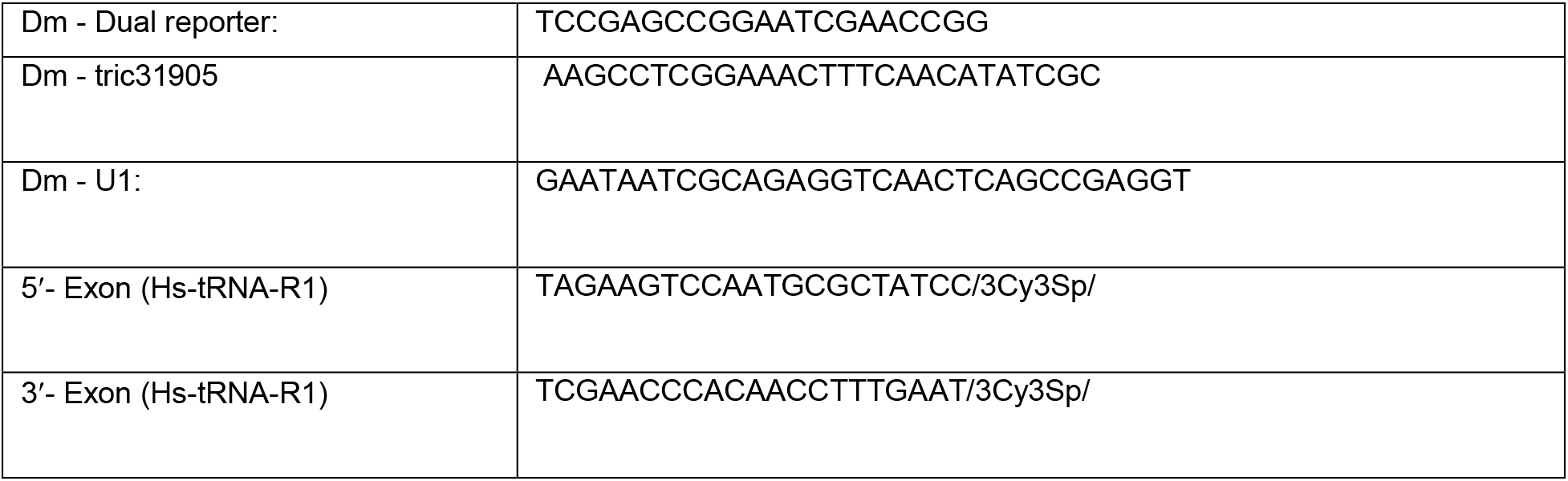
Northern blot probes

**Table S4.**
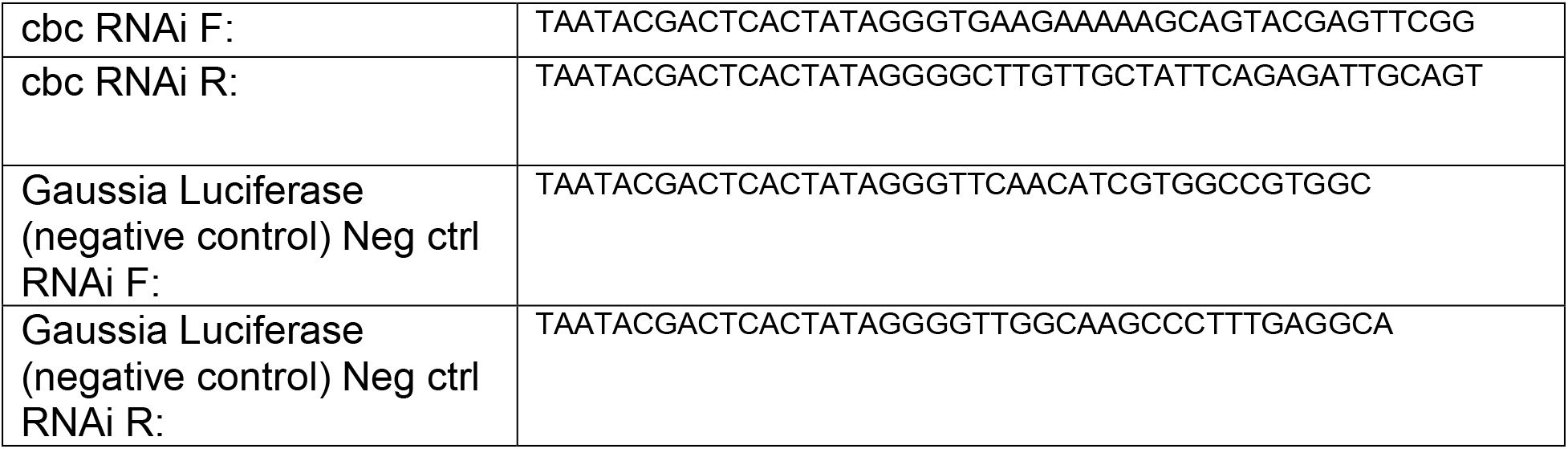
Primers for dsRNA knockdowns.

**Table S5.**
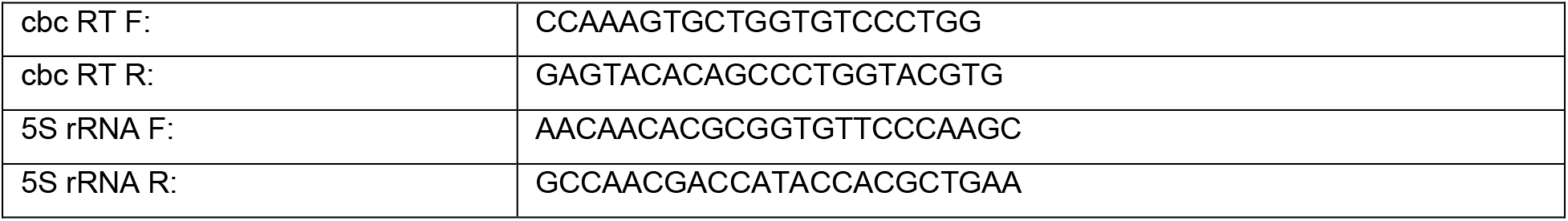
RT-PCR Primers.

